# Differential stromal reprogramming in benign and malignant naturally occurring canine mammary tumours identifies disease-promoting stromal components

**DOI:** 10.1101/783621

**Authors:** Parisa Amini, Sina Nassiri, Alexandra Malbon, Enni Markkanen

## Abstract

The importance of cancer-associated stroma (CAS) for initiation and progression of cancer is well accepted. However, as stromal changes in benign forms of naturally occurring tumours are poorly understood, it remains unclear how CAS from benign and malignant tumours compare. Spontaneous canine mammary tumours are viewed as excellent models of human mammary carcinomas (mCA). We have recently reported highly conserved stromal reprogramming between canine and human mCA based on transcriptome analysis of laser-capture-microdissected FFPE specimen. To identify stromal changes between benign and malignant mammary tumours, we have analysed CAS and matched normal stroma from 13 canine mammary adenomas and compared them to 15 canine mCA. Our analyses revealed distinct stromal reprogramming even in small benign tumours. While similarities in stromal reprogramming exist, the CAS signature clearly distinguished adenomas from mCA, suggesting that it may reliably discriminate between benign and malignant tumours. We identified strongly discriminatory genes and found strong differential enrichment in several hallmark signalling pathways between benign and malignant CAS. The distinction between CAS from adenoma and mCA was further substantiated by differential abundance in cellular composition. Finally, to determine key players in CAS reprograming between adenomas and mCA, a network-based gene screening method identified modules of co-expressing genes with distinct expression profile in benign and malignant CAS, and revealed several hub genes as potential molecular drivers in CAS. Given the relevance of canine CAS as a model for the human disease, our approach identifies potential stromal drivers of tumour malignancy with implications for human mCA.

**Summary statement:** RNAsequencing-based analysis of stromal reprogramming between benign and malignant naturally occurring canine mammary tumours identifies potential molecular drivers in cancer-associated stroma that support tumour growth and malignancy.

## Introduction

It is well accepted that the microenvironment surrounding cancer cells, the so-called cancer-associated stroma (CAS), plays a central role in both cancer initiation as well as its progression (Bissell and Hines, 2011; Hanahan and Coussens, 2012). CAS is composed of various non-cancer cells, among them fibroblasts, immune cells, vascular cells, as well as extracellular matrix. CAS affects tumour cells in several ways: directly by promoting growth and survival of tumour cells, and also by stimulating their invasive and migratory capacity, thereby promoting invasion and metastasis (Bissell and Hines, 2011; Hanahan and Coussens, 2012). Thus, CAS is considered to be a major determinant of tumour malignancy. However, in contrast to malignant tumours, it remains largely unexplored whether and what stromal changes occur in benign forms of naturally occurring cancer, and how CAS in benign neoplasms compares to that of malignant tumours in the same tissue. Such knowledge has the potential to help identify disease-promoting and/or suppressive features of CAS and identify novel prognostic and therapeutic targets therein.

Close resemblance with regards to both pathophysiology and clinical aspects have positioned spontaneously occurring tumours in the domestic dog as valuable model to enhance understanding of tumour biology in both canine and human patients (Gardner et al., 2015; Karlsson and Lindblad-Toh, 2008; Rogers, 2015). In particular, canine simple mammary carcinoma (mCA) are regarded as excellent models for human mCA, as they recapitulate the biology of human mCA both histologically and molecularly, and overcome several of the limitations of rodent tumour models (Liu et al., 2014; Queiroga et al., 2011; Schiffman and Breen, 2015). Canine simple mCA are malignant epithelial neoplasms that infiltrate the surrounding tissue, thereby inducing a strong stromal response, and can give rise to metastases (Goldschmidt et al., 2011). In contrast, canine simple mammary adenomas are well-demarcated, non-infiltrative benign mammary tumours generally associated with little fibrovascular supporting stroma (Goldschmidt et al., 2011). Whether these benign adenomas can progress into more malignant forms, such as mCA, remains an unresolved controversy.

Given the central role of CAS in human cancer in general, and mCA in particular, it is likely to also play a key role in canine mammary tumours. To understand stromal reprogramming in canine mCA and how it compares to human mCA, we have previously analysed CAS reprogramming in formalin-fixed paraffin embedded (FFPE) breast cancer tissue using by laser-capture-microdissection (LCM) and quantitative PCR (RT-qPCR), and further advanced the approach to analyse LCM subsections by next-generation sequencing (RNAseq) (Amini et al., 2017; Ettlin et al., 2017). Using this powerful RNAseq-driven approach, we have very recently assessed stromal reprogramming in a set of 15 canine mCA, and demonstrated strong molecular homology in stromal reprogramming between canine and human mCA, emphasizing the relevance of the canine model for the human disease also with regards to CAS reprogramming (Amini et al., 2019).

For diagnostic and prognostic purposes, there is a high need for markers to reliably predict the clinical course of a tumour. This necessitates understanding what differentiates benign and malignant tumours on a molecular basis. Several studies have demonstrated differences between stromal expression patterns from human mCA *in situ* compared to invasive tumours, some of which can be used as predictive markers for disease (Conklin and Keely, 2012; Yaari et al., 2013). However, data regarding stromal reprogramming in naturally occurring benign mammary tumours are inexistent. To analyse whether stromal reprogramming occurs in naturally occurring benign mammary tumours, and to compare stromal reprogramming between benign and malignant mammary tumours, we investigated 13 cases of canine mammary adenoma, and compared stromal reprogramming in canine adenoma to that in canine mCA.

## Results

### Transcriptomic profiling of matched CAS and normal stroma from canine mammary adenomas isolated by laser-capture microdissection from FFPE specimens

To characterize stromal changes associated with canine simple adenomas, we isolated both CAS and matched normal stroma (i.e. stroma adjacent to unaltered mammary glands) from 13 FFPE samples of canine simple adenoma using our established protocol (Amini et al., 2017; 2019). Patient characteristics for all adenoma cases that were included and representative images for tissue isolation can be found in Table 1 and Supplementary Figure 1. Pairwise sample-to-sample Pearson correlation analysis using all genes revealed a clear separation of normal stroma and CAS, demonstrating that CAS in adenoma also undergoes a reprogramming that clearly differentiates it from normal stroma (Figure 1A). Analysis of differentially expressed genes with a FDR cut-off of 0.05 and fold change threshold of 2 revealed 193 genes to be significantly deregulated in CAS from adenoma compared to normal stroma, including 57 significantly up- and 136 significantly down-regulated genes (Figure 1B and Supplementary Table 1). Over-representation analysis of GO terms associated with biological processes suggested changes in following GO categories: extracellular structure organisation, adhesion, response to organic substance and endogenous stimulus, regulation of multicellular organismal development, and responses related to the immune system (Figure 1C). Moreover GO terms associated with cellular components revealed main changes pertaining to the extracellular matrix (Figure 1D), which was also supported by GO terms associated with molecular functions, highlighting strong changes in binding of various ECM components (Figure 1E).

**Table 1.** Overview of canine mammary simple adenoma cases included in this study.

| Case # | Gender | Breed | Age (Years) | Simple Adenoma | Age of Sample (Months) |
| --- | --- | --- | --- | --- | --- |
| 1 | f | Rhodesian Ridgeback | 8 | yes | 13 |
| 2 | f | Deutscher Wachtelhun | 6 | yes | 20 |
| 3 | f | Golden Retriever | 8 | yes | 24 |
| 4 | f | Cocker Spaniel | 8 | yes | 22 |
| 5 | f | Coton de Tuléar | 13 | yes | 29 |
| 6 | f | Epagneul Breton | 8 | yes | 6 |
| 7 | f | Galgo Espanol | 9 | yes | 7 |
| 8 | f | Pinscher | 11 | yes | 15 |
| 9 | f | Basset | 12 | yes | 9 |
| 10 | f | Chihuahua | 9 | yes | 8 |
| 11 | f | Yorkshire Terrier | 10 | yes | 11 |
| 12 | f | Mittelpudel | 11 | yes | 4 |
| 13 | f | West Highland White Terrier | 9 | yes | 4 |

Clinical data from dogs with simple mammary adenoma; Case # = case number as referred to within this study; f/n = female, neutered; n.d. = not disclosed; age = age at excision of tumour; age of sample = time between initial tumour excision and sampling of stroma/RNA extraction.

**Figure 1:**
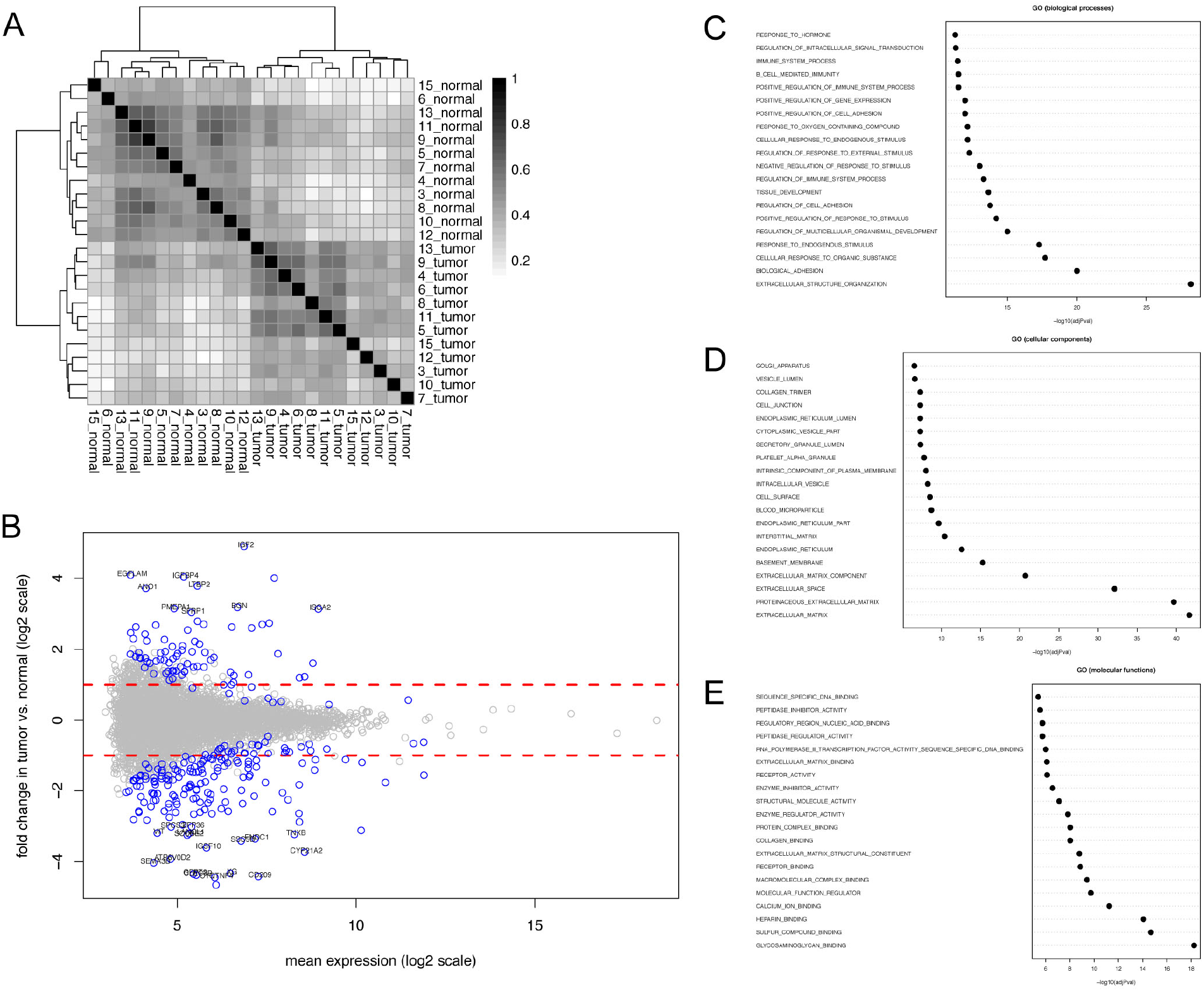
RNAseq-based transcriptomic analysis of cancer-associated stroma and matched normal stroma from 13 canine simple mammary adenoma. **A)** Pairwise Pearson correlation analysis of cancer-associated stroma (CAS) and normal stroma samples isolated from canine simple adenoma. The analysis was performed using all genes. **B)** Scatter plot of fold change versus mean expression highlighting differentially expressed genes in tumour stroma compared to normal stroma, using FC>2 and FDR<0.05 as cut-off values. **C)-E)** Top 20 over-represented Gene Ontology (GO) terms associated with biological processes **(C)**, cellular components **(D)**, and molecular functions **(E)** among genes significantly de-regulated in CAS compared to normal stroma.

Validation of RNAseq data was achieved through RT-qPCR of 8 strongly up- and down-regulated genes (SCUBE2, MMP2, VIT, SDK1, STRA6, IGF2, PIGR, and SFRP1), all of which showed significant expression changes consistent with RNAseq (Figure 2A-I). Up-regulation of α-smooth muscle actin (α-SMA) in CAS compared to normal stroma was further evident on protein level by immunofluorescence (IF), in line with RNAseq results (ACTA2 log2 fold-change 0.875, p-value 0.0015; Figure 2K). Similarly, vimentin expression decreased from normal stroma to CAS, consistent with sequencing results (VIM log2 fold-change −1.044, p-value 2.58 E-08; Figure 2L). Taken together, our data clearly demonstrate the occurrence of extensive stromal reprogramming in these benign naturally occurring tumours that is mainly driven by changes in the extracellular matrix, fibroblast activation and components of the immune system.

**Figure 2:**
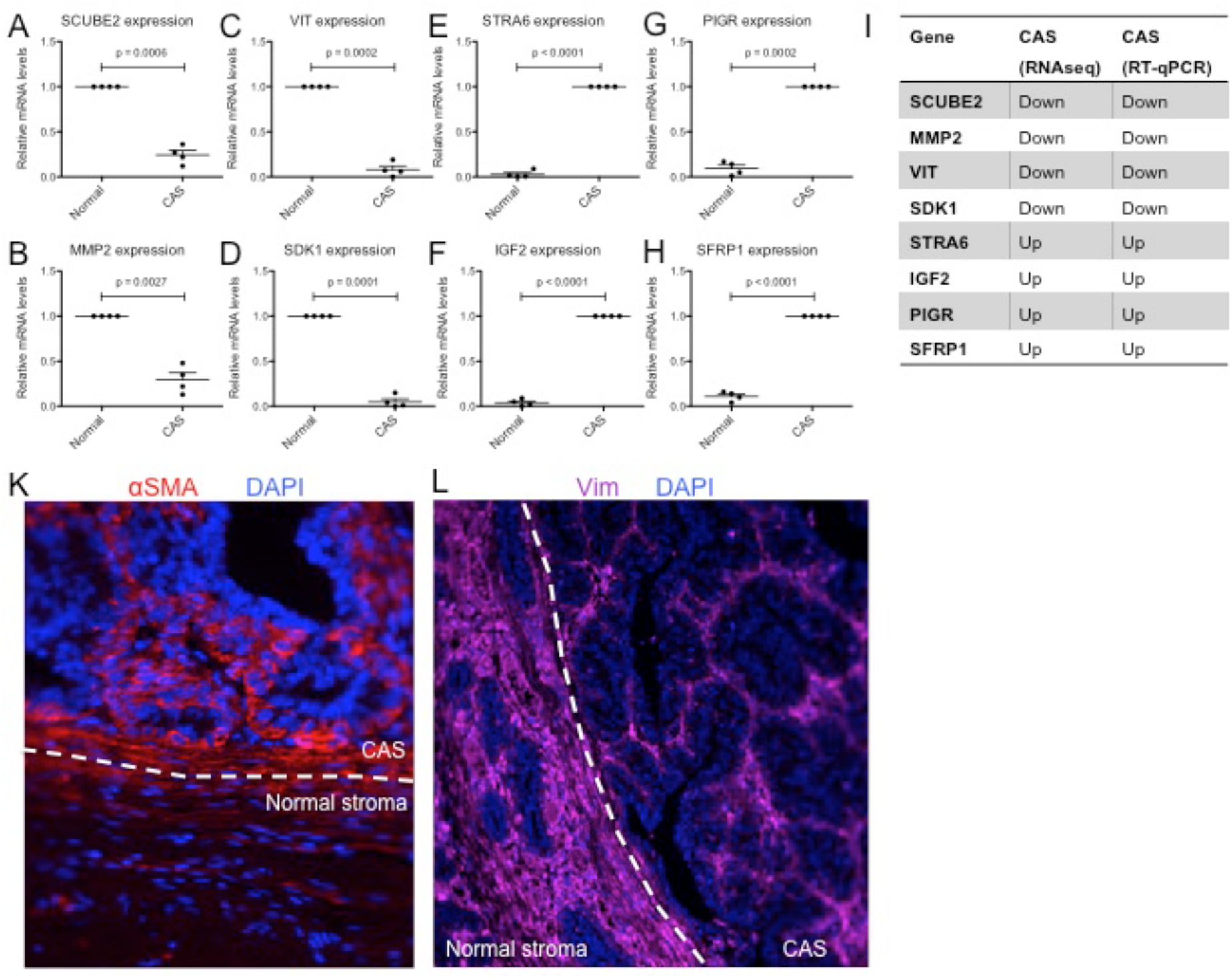
Validation of selected genes from the adenoma data by IF and RT-qPCR. **A)-H)** Relative mRNA levels of CAS-associated genes in normal stroma and CAS isolated by laser-capture microdissection, measured by RT-qPCR. **(A)**: SCUBE2; **(B)**: MMP2; **(C)**: VIT; **(D)**: SDK1; **(E)**: STRA6; **(F)**: IGF2; **(G)**: PIGR; **(H)**: SFRP1; Values are mean values ±SEM of four independent cases, normalized to expression levels in normal stroma (for SCUBE2, MMP2, VIT, and SDK1), or CAS (STRA6, IGF2, PIGR, and SFRP1), respectively. P-values were calculated using student’s t-test, and significance cutoff was set at p = 0.05. **(I)** Summary of the expression trends as detected by RT-qPCR and RNAseq. **(K and L)** Immunofluorescent staining of α-SMA (red) **(K)**, or Vimentin (purple) **(L)** in CAS and normal stroma of a representative canine simple mammary adenoma. Dapi staining (blue) visualizes cell nuclei. The dashed white line indicates the border between CAS and normal stroma.

### The CAS signature distinguishes benign from malignant mammary tumours

To understand how stromal reprogramming in benign canine mammary adenomas compares to malignant mCA, we juxtaposed our dataset for CAS in adenoma to a dataset of matched CAS and normal stroma from 15 canine mCA that we had obtained using the same methodology as previously reported (Amini et al., 2019). As the histopathological appearance of normal, uninvolved stroma showed no difference between adenomas and mCA as expected, we merged the two data sets while adjusting for potential batch effects (see methods for details). Interestingly, PCA of the combined data showed three homogenous yet distinct clusters with the first two principal components clearly separating CAS from adenoma and mCA from each other as well as from their normal counterparts, supporting the notion that stromal reprogramming is strongly influenced by the type of tumour (Figure 3A). Furthermore, adenoma-derived stroma seemed to be much more similar to normal stroma than CAS from mCA, suggesting that CAS undergoes a gradual change during the development of malignant tumors.

**Figure 3:**
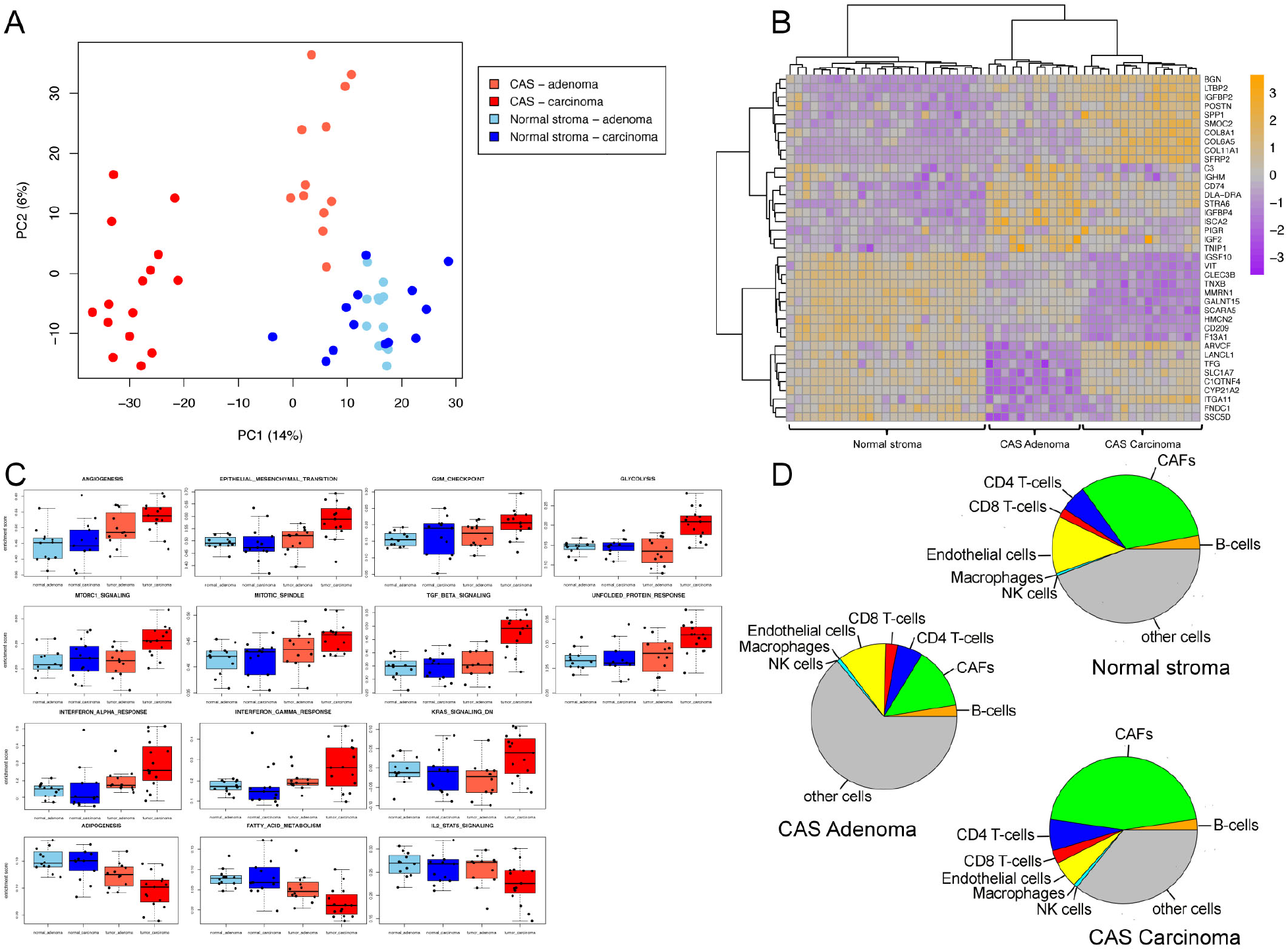
The cancer-associated stromal signature distinguishes benign from malignant canine mammary tumours. **A)** Principal component analysis (PCA) of the batch-adjusted adenoma and mCA data combined. Defining each study as one batch, combined data was adjusted for potential batch effects under the assumption that normal stroma is similar between adenoma and mCA. Dark red = CAS in mCA, bright red = CAS in adenoma, dark blue = normal stroma from mCA cases, bright blue = normal stroma from adenoma cases. B) Heatmap of 40 most discriminatory genes between CAS in adenoma, CAS in mCA and normal stroma from both conditions, as revealed by PLSDA. Top 20 features with largest absolute loading were selected separately for the first and second components of the PLSDA model. Full list of PLSDA loadings can be found in Supplementary Table 2. C) single sample GSEA analysis of hallmark pathways (obtained from MSigDB) in normal stroma and CAS from both adenoma and mCA. Pathways with an ANOVA p-value smaller than 0.05 are shown. D) Pie charts summarising the average cellular composition of CAS as obtained from the EPIC algorithm. Pie area corresponds to average cellular fraction in the respective group.

We next used Partial Least Squares Discriminant Analysis (PLSDA) to identify characteristic features of each cluster and assess the contribution of individual genes to the observed separations. Intuitively, PLSDA can be viewed as a supervised extension of PCA, in which principal components are rotated such that maximum separation is achieved between groups of observations. The loading vector of the PLS components can then be used to extract most discriminatory features. The expression profile of 40 most discriminatory genes as revealed by PLS loadings (Figure 3B, Supplementary Figure 2 for seleted genes, and Supplementary Table 2 for the full list of genes) revealed several interesting clusters of genes: i) genes that are strongly up-regulated in CAS from mCA, but remain practically unchanged in CAS from adenoma compared to normal stroma (IGFBP2, POSTN, COL11A1, SFRP2, and others); ii) genes whose expression is similar between CAS from mCA and normal stroma, but strongly increases in CAS from adenoma (such as CD74, STRA6, and PIGR); iii) genes that are strongly down-regulated in CAS from mCA (e.g. IGSF10, HMCN2, and CLEC3B); and iv) and genes that are strongly down-regulated specifically in CAS from adenoma (e.g. ARVCF, LANCL1, ITGA11, etc.). These genes are potentially interesting targets associated with observed differences in benign versus malignant tumours. For instance, ARVCF is a member of the catenin protein family, best known for their role in cell-cell adhesion and in canonical Wnt signalling at the heart of epithelial to mesenchymal transition (Kourtidis et al., 2017; Zhang et al., 2015). As validation of the PLSDA findings, we also performed IHC for ARVCF on mCAs and adenomas. In agreement with the RNAseq data, while moderate ARVCF staining could be detected in normal stroma of adenomas, its intensity strongly decreased in CAS from adenoma (Supplemetary Figure 3A); whereas it even increased from normal stroma to CAS in mCA (Supplemetary Figure 3B). The mechanistic relevance of ARVCF deregulation and the other identified identified targets to CAS reprogramming during tumour growth and progression should be interrogated in future studies.

mCA differ in clinical and molecular aspects from adenomas. It is therefore reasonable to assume that distinct tumour-promoting pathways may be at play in CAS from mCA compared to that in adenomas. Indeed, enrichment analysis of hallmark pathways among CAS from mCA, CAS from adenoma, and normal stroma revealed several hits. Angiogenesis, epithelial-to-mesenchymal transition, G2M checkpoint, glycolysis, interferon alpha and gamma responses, KRAS signalling down, mitotic spindle, mTORC1 signalling, TGFbeta signalling and unfolded protein response showed increased enrichment in CAS from mCA, whereas adipogenesis, fatty acid metabolism and IL2-STAT5 signalling displayed a significantly decreased enrichment between CAS from mCA compared to the other groups (Figure 3C, Supplementary Figure 4). Interestingly, for many of these pathways in adenoma CAS showed an intermediate enrichment between normal stroma and CAS from mCA, supporting the notion of progressive stromal adjustments to malignant transformation of the associated epithelium.

To further assess the contribution of changes in cellular composition to the observed transcriptional reprograming of CAS in adenoma versus mCA, we utilized a previously established algorithm to estimate the proportion of main immune and stromal cells from CAS gene expression data (Racle et al., 2017). Of the cell types quantified, cancer-associated fibroblasts (CAFs) and endothelial cells made up a large portion of the cellular composition of CAS in all groups (Figure 3D, Supplementary Figure 5). Of note, detection of CAFs in normal stroma most probably reflects the inherent difficulties in differentiating between fibroblasts and CAFs, and as such the CAFs detected by this methodology can be interpreted as ‘fibroblast-like cells’ either in normal stroma or cancer stroma. We found the fraction of CAFs to be higher in CAS from mCA versus adenomas. In contrast, the relative abundance of endothelial cells was lower in CAS from mCA compared to CAS from adenoma. These findings suggest that the CAS reprograming as manifested in deregulation of genes and pathways is, at least in part, influenced by changes in the cellular composition of the stroma.

Taken together, our data demonstrate that CAS signature clearly distinguishes benign adenomas from malignant canine mCA, suggesting CAS could be a discriminatory feature influencing the clinical course of the disease. Furthermore, our analyses identify novel stromal targets associated with tumour malignancy, and reveal specific perturbations of several candidate genes and signalling pathways as well as cell types to be associated with CAS of malignant tumours.

### Coexpression network analysis reveals modules and hub genes associated with CAS obtained from adenomas and mCA

Complementary to differential expression and gene set enrichment analysis which focus on individual genes and previously annotated gene sets, weighted gene co-expression network analysis (WGCNA) is a systems approach that allows for unbiased screening of genes based on their interconnectedness, thus revealing the inherent organization of the transcriptome that underlies the biology of interest and pointing out candidate targets and biomarkers for further investigation (van Dam et al., 2017; Zhao et al., 2010). To extend our understanding of the CAS transcriptional reprogramming in adenoma and mCA, we applied WGCNA on a subset of highly variable genes, and identified six clusters of highly positively correlated genes, hereafter referred to as gene modules (Figure 4A, Supplementary Figure 6, Supplementary Table 3). Closer inspection of module eigengenes as the summary of the expression pattern within each module revealed three potentially interesting modules: module turquoise showed overexpression in CAS from adenomas, whereas modules brown and yellow showed overexpression in CAS from mCA (Figure 4B). In agreement with our earlier findings, the observation that distinct modules were associated with CAS from different origin further highlights differential transcriptional reprograming between CAS obtained from adenomas and mCA. Finally, we examined intramodular connectivity to identify hub genes within each module of interest (Supplementary Table 3). Figure 4C provides network visualization of selected modules and highlights top 5 genes with largest intramodular connectivity score serving as candidate hub genes for each module. For the turquoise module, these were SPINK5, DSG1, FLG2, KRT1, and DMKN; for module yellow the top 5 candidates consisted of CDH1, ST14, EHF, KRT8, and KIAA1217; and for the module brown we identified COL8A2, SORCS2, BGN, RUNX1, and IGFBP2 as the top 5 genes. Collectively, these findings reveal the inherent structure of the transcriptome underlying CAS and its modular distinction between adenoma and mCA. The identified hub genes serve as potential biomarkers or candidate targets for pharmaceutical intervention and will be subject of future studies.

**Figure 4:**
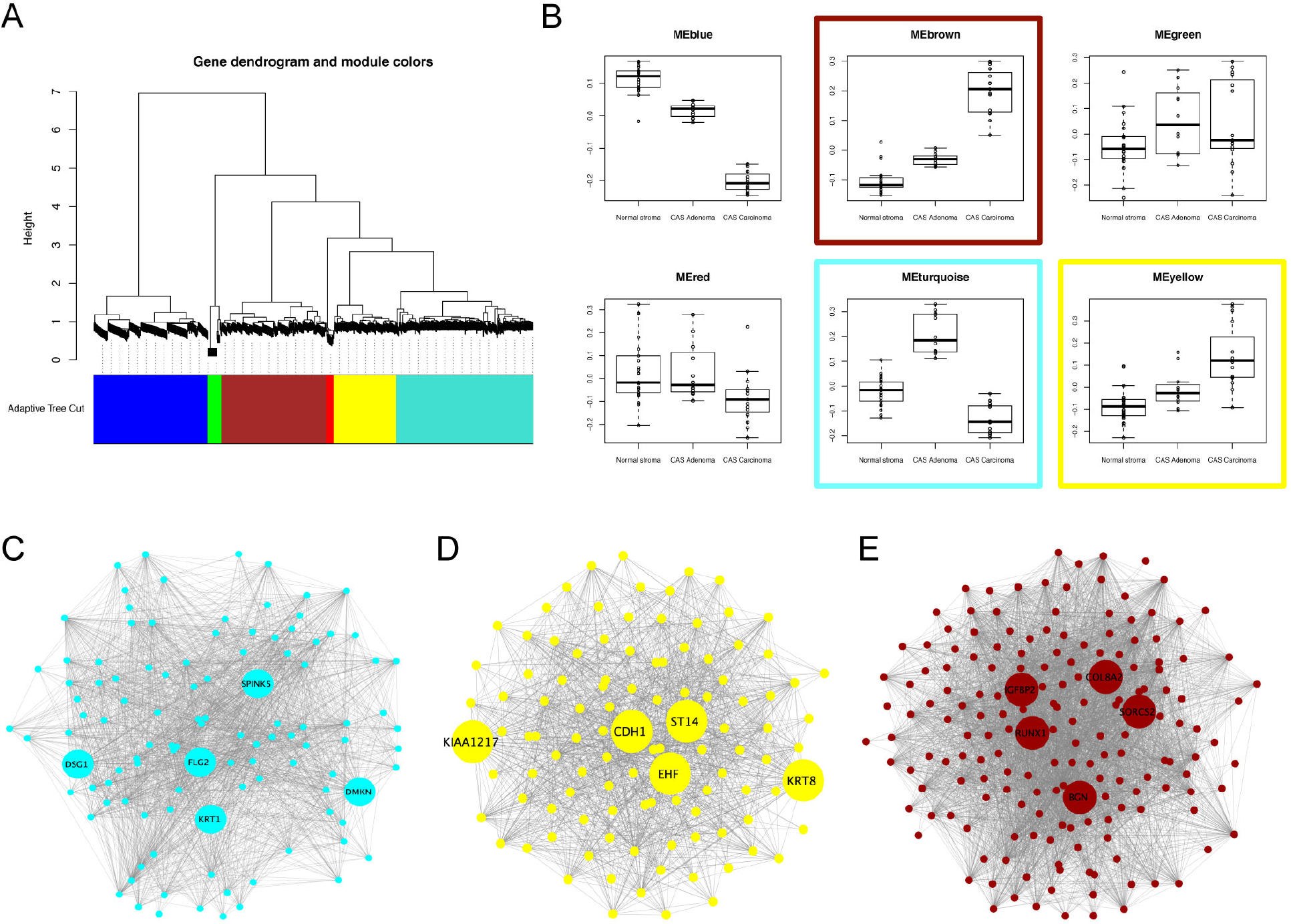
Weighted gene coexpression network analysis reveals modules and hub genes associated with CAS from adenomas and mCA. A) Modules of highly positively correlated genes withig the top 10% highly variable genes. B) Association between module eigengenes and biological groups suggests three modules of interest: turquoise, brown, and yellow. C) Network visualization of selected modules and their respective hub genes.

## Discussion

CAS plays a key role in cancer initiation and progression in human cancer (Bissell and Hines, 2011; Hanahan and Coussens, 2012). For diagnostic and therapeutic purposes, there is significant interest in understanding how CAS differs between benign and malignant forms of the disease. This necessitates understanding what differentiates mCA from adenomas on a molecular basis. Stromal gene expression has indeed been shown to predict clinical outcome in invasive breast cancer (Calvo et al., 2013; Finak et al., 2008). Several studies have demonstrated differences between stromal expression patterns from human tumours *in situ* compared to those that display invasive properties, some of which can be used as predictive markers for disease (e.g. (Conklin and Keely, 2012; Ma et al., 2009; Yaari et al., 2013)). Similarly, studies using mouse models have analysed changes in stromal cell populations at different stages of mCA progression (e.g. (Calvo et al., 2013)). To the best of our knowledge, however, there is no dataset (human or other) available that describes stromal reactions in naturally occurring benign mammary adenomas that could be used to compare to malignant mCA. Thus it remains unknown whether the stroma around benign adenomas undergoes reprogramming, and if so, what changes occur at the molecular level. As a consequence of this, it is unclear how these changes in benign neoplasms compare to malignant tumours of the same tissue. Such knowledge has the potential to help identify both disease-promoting and/or suppressing features of CAS and identify novel prognostic and therapeutic targets therein. Given that canine mammary tumours are regarded as valuable model for human breast cancer, and the central role of CAS in human cancer in mCA, we have recently analysed stromal reprogramming in canine mCA, and demonstrated the presence of strong molecular homology in stromal reprogramming between canine and human mCA, emphasizing the relevance of the canine model for the human disease (Amini et al., 2017; 2019; Ettlin et al., 2017). To understand whether stromal reprogramming also occurs in benign mammary tumours, and to compare stromal reprogramming between benign and malignant mammary tumours, we have now analysed stromal reprogramming in 13 cases of canine mammary adenoma and compared it to that in canine mCA.

Here we report that CAS in canine adenoma undergoes a reprogramming that clearly differentiates it from normal stroma, with major changes in GO terms related to extracellular structure organisation, adhesion, response to organic substance and endogenous stimulus, regulation of multicellular organismal development, immune responses, and the extracellular matrix (Figure 1). These changes are consistent with fibroblast- and immune-cell driven remodelling of the tumour microenvironment, which are known to be heavily involved in tumour biology of mCA (Bissell and Hines, 2011; Hanahan and Coussens, 2012). Indeed, we find clear signs of fibroblast activation and reprogramming, as evidenced by the increase in αSMA and the decrease in vimentin by IF (Figure 2), which is also seen in malignant canine mCA (Amini et al., 2019; Ettlin et al., 2017; Yoshimura et al., 2011). It is interesting that the stroma surrounding adenomas shows such a clear reprogramming, as these non-infiltrative benign mammary tumours are generally associated with little fibrovascular supporting stroma (Goldschmidt et al., 2011). Of note, αSMA reactive fibroblasts or slight upregulation of several matrix-metalloproteinases and their inhibitors have been detected in benign and malignant lesions of the human breast (Barth et al., 2001; Brummer et al., 1999; Elenbaas and Weinberg, 2001; Sappino et al., 1988; Surowiak et al., 2007; Yamashita et al., 2012), as well as the canine mammary gland (Yoshimura et al., 2011). Hence, this first detailed glimpse into CAS surrounding spontaneous benign tumours of the breast suggests that stromal reprogramming is an early reaction to development of benign tumours and is characterized by strong transcriptional responses, the molecular minutiae of which have to be elucidated in future studies. The obvious stromal reprogramming adjacent to adenomas suggests a subset of alterations in the extracellular matrix to be early events in reactive stroma during initial tumour development, and not depend on tumour malignancy. Fibroblasts are the most abundant cells of the connective tissue, and responsible for production and maintenance the extracellular matrix (Chen and Song, 2019). Due to a very strong innate plasticity of fibroblasts, these cells are highly reactive towards changes in their environment, which gives rise to their inherent heterogeneity with regards to both phenotype and function. This makes them ideally adapted to fulfil very diverse roles ranging from maintenance of physical tissue support, wound healing to modulating inflammatory processes and supporting tumour growth. It thus seems that these changes in extracellular matrix composition might be mainly driven by fibroblast activation. Of note, this activation does not necessarily equal an increase in numbers of fibroblasts present, but may simply reflect their transcriptional status. It is well accepted, that the composition of the extracellular matrix strongly changes during tumour progression (Bissell and Hines, 2011). Recent advances have revealed a multitude of specific subpopulations of cancer-associated fibroblasts that display strong phenotypic diversity and functional heterogeneity (e.g. (Chen and Song, 2019; Costa et al., 2018)). It will be of great interest now to investigate the detailed changes that discriminate the ‘early fibroblastic reaction’ that might be indiscriminate towards hyperplasia within a given epithelium from the reaction associated with malignant tumours to understand how the fibroblastic response changes in relation to tumour cell malignancy, and vice versa.

By comparing stromal reprogramming in benign canine mammary adenomas to malignant mCA, we identified a list of targets that clearly distinguish CAS of benign adenomas from that in malignant mCA, further supporting the notion CAS has the potential of being a discriminatory feature influencing the clinical course of the disease (Figure 3). Further, we found specific perturbations of e.g. EMT and glycolysis to be associated with CAS of malignant tumours. EMT-related genes such as COL11A1, COL8A2, and ADAM12 that are overexpressed in mCA could thus be potential biomarkers for canine invasive mCA, similarly to human mCA (Freire et al., 2015; Kleinert et al., 2015; Ma et al., 2015). In the glycolysis pathway, PLOD1/2, FUT8 and TSTA3 are deregulated genes that participate in metabolism and glycolytic processes, which can influence the malignant transformation of cells, tumour development and metastasis (Gilkes et al., 2013; Kim et al., 2012; Tu et al., 2017). Among the targets that are strongly differentially expressed between CAS from mCA and adenoma, we validated the selective down-regulation of ARVCF in adenoma. ARVCF is a delta catenin family member involved in protein-protein interactions at adherens junctions (Lelièvre, 2010). Increased amounts of ARVCF have been suggested to disrupt cell adhesion (Reintsch et al., 2008), suggesting a possible role in cancer progression (Zhang et al., 2015). However, to date, there is nothing specific known about a possible role of ARVCF in CAS biology. Further functional assessment of ARVCF as well as the other differentially expressed targets and their association with tumour malignancy should be determined in future studies.

While not completely resolved, there is clear evidence that canine mammary gland tumours are a continuum from benign to malignant, supporting the comparison of canine mammary adenomas and mCA as different states of malignancy of the same disease (Sorenmo et al., 2009). Interestingly, PCA of the combined adenoma and carcinoma CAS shows adenoma-derived stroma to be much more similar to normal stroma than CAS from mCA, suggesting that CAS undergoes a gradual change during the development of malignant tumors. This was further corroborated by both PLS-DA and enrichment analysis of hallmark pathways among CAS from mCA, CAS from adenoma, and normal stroma, where many of these pathways in adenoma CAS showed an intermediate enrichment between normal stroma and CAS from mCA (Figure 3). With respect to the cellular composition of CAS, cancer-associated fibroblasts (CAFs) and endothelial cells made up a large portion of the cellular composition of CAS in all groups. We found the fraction of CAFs to be higher in CAS from mCA versus adenomas, suggesting CAFs to be strong drivers towards tumour malignancy, consistent with current literature (Chen and Song, 2019). The relatively high number of CAFs in the normal stroma likely reflects the difficulties in differentiating between normal fibroblasts and CAFs based on subtle differences in their gene expression profiles. The lower relative abundance of endothelial cells in CAS from mCA compared to CAS from adenoma is in line with the observation that malignant tumours often harbour large hypoxic or even necrotic areas due to insufficient vascular supply in relation to their strong proliferative properties. Indeed, a hypoxic tumour microenvironment and tumour progression are strongly linked (Petrova et al., 2018). Thus, we find that the CAS reprograming as manifested in deregulation of genes and pathways is, at least in part, also influenced by changes in the cellular composition of the stroma. Future work exploiting CAS composition e.g. using single cell methods to simultaneously characterise changes in composition and state of CAS at the single cell resolution between benign and malignant tumours would help elucidating such changes further.

Identification of the hub genes (Figure 4) SPINK5, DSG1, FLG2, KRT1, and DMKN the turquoise module, which is strongly decreased in CAS from mCA compared to adenoma, is highly interesting, since all of these genes have been strongly linked to maintenance of epithelial differentiation and integrity (Deraison et al., 2007; Hammers and Stanley, 2013; Leclerc et al., 2014; Mohamad et al., 2018; Roth et al., 2013). This suggests an important function of stromal reprogramming in destabilization of epithelial differentiation and integrity. For module yellow, which is progressively up-regulated from normal stroma to adenoma to mCA, the top candidate hub genes consisted of CDH1, ST14, EHF, KRT8, and KIAA1217, all of which have important roles in epithelial cells, and/or are associated with tumour malignancy (Lee et al., 2016; Moll et al., 2008; Uhland, 2006; Yamazaki et al., 2015). Similarly, hub genes of module brown, COL8A2, SORCS2, BGN, RUNX1, and IGFBP2, displayed a progressive increase from normal stroma to CAS in adenoma and mCA. These genes have important functions in the extracellular matrix and cell differentiation, and some have been associated with tumor progression or bad outcome in breast cancer (Finak et al., 2008; Schaefer et al., 2016; Scheitz and Tumbar, 2013; VanOudenhove et al., 2016; Yau et al., 2015). Collectively, these findings reveal differences in transcriptional reprogramming in CAS between benign and malignant breast tumours, and identify hub genes that could serve as potential biomarkers or candidate targets for pharmaceutical intervention. The detailed elucidation of the impact of these candidates to disease progression shall be subject of future studies.

To conclude, we provide a first detailed view of stromal reprogramming in naturally occurring benign mammary adenomas, which demonstrates the occurrence of strong stromal reprogramming even in small benign tumours. Furthermore the CAS signature clearly distinguishes benign adenomas from malignant mCA, allowing identification of several hub genes as potential molecular drivers in CAS. Given the relevance of canine CAS as a model for the human disease, our approach identifies potential stromal drivers of tumour malignancy with implications for human mCA.

## Materials and methods

### Aim, case selection and tissue processing

We isolated CAS and matched normal stroma from FFPE tissue sections of canine mammary adenoma by LCM for transcriptome analysis by RNAseq. For this, thirteen canine simple mammary adenoma samples were obtained from the Institute of Veterinary Pathology of the Vetsuisse Faculty Zürich (Table 1). All samples were archival formalin-fixed, paraffin-embedded tissue samples either from the Animal Hospital of Zurich or external referral cases from veterinarians practicing in Switzerland. Details regarding selection criteria are described in (Ettlin et al., 2017). Paraffin blocks were routinely kept at room temperature. Tissue processing for LCM was performed as described in (Amini et al., 2017). All cases were reviewed by a veterinary pathologist. Criteria for case selection included female dogs, simple mammary adenoma, and sufficient tumour stroma content for tissue isolation. Table 1 provides clinical details, such as age and breed of each patient, sample age and tumour type, for all cases included in the study.

### Laser-capture microdissection

Laser capture microdissection (LCM) for selective isolation of matched CAS and normal stroma was performed as previously described ((Amini et al., 2017; 2019; Ettlin et al., 2017)). Areas for dissection were reviewed by a veterinary pathologist. Highly enriched populations of normal or tumour-associated stroma were identified and isolated according to the manufacturer’s protocol. Normal stroma samples were isolated from the same slides, from regions specified by a pathologist that were adjacent to unaltered mammary glands and presented no obvious alterations and were at least 2-4 mm away from the tumour, in accordance with established procedures (Finak et al., 2008). Isolation of cells of interest was verified by microscopic examination of the LCM cap as well as the excised region after microdissection (Supplementary Figure 1).

### RNA isolation

RNA was isolated using the Covaris® truXTRAC FFPE RNA kit and the Covaris® E220 focused ultrasonicator as described in (Amini et al., 2017). Details about RNA concentration, yield, and quality for adenoma samples can be found in Supplementary table 4.

### Quantitative RT-PCR

Quantitative RT-PCR using Taqman® primers was performed as described in (Amini et al., 2017). Primers are detailed in Supplementary table 5.

### Immunofluoresence

Immunofluorescence was performed as outlined in (Amini et al., 2019). Antibodies and conditions used for immunofluorescence are detailed in Supplementary table 6.

### RNA sequencing library preparation

10 ng of RNA from Elution 1 (E1) diluted to a concentration of 0.33 ng/μl in a total volume of 30 μl was submitted for next-generation RNA sequencing and analysed as outlined in (Amini et al., 2019).

### Bioinformatics analyses

RNAseq quantification was performed with kallisto 0.44.0 with sequence-based bias correction using transcript sequences obtained from ENSEMBLE (CanFam3.1) (Bray et al., 2016). Kallisto’s transcript-level estimates were further summarized at the gene-level using tximport 1.8.0 from Bioconductor (Soneson et al., 2015). Both raw data and gene-by-sample matrix of estimated counts have been deposited online and are publicly accessible from Gene Expression Omnibus (GEO) under accession number GSE135454. One of the normal stroma samples (14_normal) had extremely low sequencing depth and therefore was excluded along with its CAS pair (14_tumor) from downstream analyses. Carcinoma data was processed similarly as previously reported (Amini et al., 2019), and can be accessed from GEO under GSE135183.

Prior to downstream analyses, lowly abundant genes were filtered out, and except for differential expression analysis, mean-variance trend was adjusted for using the variance-stabilizing transformation from DESeq2 1.22.0 package (Love et al., 2014). Pairwise sample Pearson correlation was computed in the adenoma data with top 10% most variable genes, and visualized using the pheatmap R Package (Kolde), with clustering distance and method set to Euclidean and ward.D2, respectively. Differentially expressed genes were identified using DESeq2 1.22.0 (Love et al., 2014), with FDR=0.05 and FoldChange=2 as significance thresholds. Over-representation analysis of Gene Ontology terms among significant genes was performed using the MSigDB webtool (http://software.broadinstitute.org/gsea).

For comparisons involving adenoma and carcinoma samples, the two datasets were merged using an intersection of genes present in both, followed by removal of lowly abundant genes. Treating each study as one batch and under the assumption that normal stroma is similar between adenoma and mCA, merged expression data was adjusted for potential batch effects using the ComBat empirical Bayes framework as implemented in the SVA 3.30.1 from Bioconductor (Leek et al.). To further mitigate technical noise, batch-corrected expression data was further adjusted for global differences across samples using quantile normalization as implemented in the limma 3.38.3 package from Bioconductor (Ritchie et al., 2015). PLSDA was implemented using mixOmics 6.6.2 from Bioconductor (Rohart et al., 2017), with the number of components included in the model set to 2. Single-sample gene set enrichment analysis was performed using the ssgsea functionality within GSVA 1.30.0 from Bioconductor (Hänzelmann et al., 2013). In silico enumeration of cell types from bulk tissue gene expression data was performed using the EPIC algorithm (Racle et al., 2017).

Coexpression network analysis was performed using WGCNA R package (Langfelder and Horvath, 2008). In brief, pairwise similarities were computed between top 10% most variable genes using biweight midcorrelation, and converted to an adjacency matrix with soft thresholding power set to 8. A signed topological dissimilarity matrix was then computed based on the adjacency matrix, and hierarchically clustered using the Ward’s minimum variance method. Following adaptive branch pruning of the clustering dendrogram, six gene modules were identified. To summarize the expression pattern within each module, module eigengene (defined as the 1^st^ principal component) was computed, and aligned along the average expression of the module to enhance interpretability. Finally, hub genes were identified based on the connectivity of nodes to other nodes within the same module, and visualized using Cytoscape (Shannon et al., 2003).

## Acknowledgements

The authors thank the histology laboratory of the Institute of Veterinary Pathology, University of Zürich for slide preparation and technical assistance, as well as Dr. Maria Domenica Moccia and Dr. Lennart Opitz (Functional Genomics Center Zürich) for their expertise regarding next-generation RNA sequencing.

## Data availability

Raw and processed sequencing data reported in this study have been deposited to Gene Expression Omnibus with the primary accession number GSE135454 and GSE135183. All other data supporting our findings is contained in the manuscript, in Supplementary figures 1 – 6 and Supplementary tables 1-6.

## Authors contributions statement

PA performed LCM, isolation of RNA, RT-qPCR, and data analysis. SN was responsible for data processing and bioinformatics analyses. AM is a board-certified veterinary pathologist, and performed and supervised choice of clinical cases and selection of areas of interest by LCM. EM was responsible for study design, supervision, data analysis and funding. PA, SN and EM wrote the first draft of the manuscript. All authors read, contributed to, and approved the final manuscript.

## Ethics approval and consent to participate

No animals were killed for the purpose of this research project, as the tissue analysed had been surgically removed in a curative setting with the verbal consent of the patient owners. According to the Swiss Animal Welfare Law Art. 3 c, Abs. 4 the preparation of tissues in the context of agricultural production, diagnostic or curative operations on the animal or for determining the health status of animal populations is not considered an animal experiment and, thus, does not require an animal experimentation license. All the used FFPE specimen were obtained for diagnostic reasons and do therefore not require a formal ethics approval, in full compliance with national guidelines.

